# An Oropouche virus *N*-glycosylation mutation in the Gc protein head domain is associated with transmissible infection in *Anopheles* mosquitoes

**DOI:** 10.64898/2026.11.23.768691

**Authors:** Luis Alonso-Palomares, John P. Collins, Bright Agbodzi, Xiang-Jun Lu, Danmeng Li, Thomas Briese, John A. Lednicky, Natasha L. Tilston, J. Kenneth Wickiser, Rhoel R. Dinglasan

## Abstract

The Oropouche virus (OROV) outbreak in South America raises concern about spreading to new foci. Although biting midges are primary OROV vectors in the Americas, we tested whether mutation and bottleneck selection could enable mosquito transmission. We used OROV240023 cultured in C6/36 cells to orally infect mosquito species. Infectious virus was detected in *Aedes aegypti* and *Anopheles gambiae* saliva, with the latter demonstrating faster dissemination at 7 days post-infection. Direct sequencing of virus from mosquito saliva identified a consensus nonsynonymous K614N substitution and a potential *N*-glycosylation site in the Gc glycoprotein involved in cell attachment. Bi-glycosylation of Gc has been reported for orthobunyaviruses infecting *An. gambiae* in East Africa. The K614N variant was a minor population (0.8%) in the input mosaic virus population used to infect mosquitoes. Virus isolated from mosquito saliva and subsequent C6/36 culture showed selection of the K614N variant. While the mechanism needs further elucidation, these findings suggest mosquito infection is possible, especially where vector populations are at higher density. Furthermore, the data warrant reconsideration of OROV240023 transmission risk beyond the Americas.

## INTRODUCTION

The 2022-2024 outbreak of Oropouche virus (OROV) in South America, with more than 10,000 confirmed cases in both endemic and non-endemic regions of Bolivia, Brazil, Colombia, Cuba, Panama, and Peru, resulted in several fatalities in healthy adults as well as scores of suspected transmissions from infected pregnant women to the fetus and associated adverse birth outcomes [1]. OROV infections have also been confirmed in 94 febrile U.S. travelers from Cuba. Of these, 14 patients were hospitalized, and 1 death remains tentatively attributed to OROV [2]. Studies indicate that biting midges, not mosquitoes, are the primary arthropod vectors responsible for OROV transmission [3-5]. Herein, we explored this assessment by determining whether mutation and bottleneck selection could enable OROV transmission by mosquitoes.

## RESULTS AND DISCUSSION

OROV from a Floridian traveler returning from Cuba on July 13, 2024 (OROV240023; GenBank accession numbers PQ064919.1, PQ064920.1, and PQ064921.1) [3] was grown in Vero E6, Aag2, and C6/36 cells, and virus was isolated from the culture supernatant and sequenced to ascertain whether cell culture passage and cell type can influence OROV genotype. We observed that the consensus sequence across all three cell cultures was remarkably similar (Supplemental Table S1) and did not differ significantly from the published genome for this clinical isolate [3]. Maximum-likelihood phylogenetic analysis revealed that C6/36-derived OROV (C1-UF) clustered with 2023–2024 OROV strains from Brazil, Colombia, Peru, and Cuba, with 100% bootstrap support (Supplemental Fig 1-3). The L segment showed the highest nucleotide identity (99.93%) to two strains isolated in Peru in 2024 (PP966984 and PP966986) and was classified as genotype IV (Supplemental Fig 1). The M segment of C1-UF shared the highest nucleotide identity (99.95%) with strain PQ363069, imported from Cuba to Italy in 2024, and was classified as genotype V (Supplemental Fig 2). The S segment displayed the closest nucleotide identity (99.79%) to four strains deposited in GenBank (PP477311, PQ363070, PP966968, and PP952117), originating from Colombia, Peru, and two Cuban cases imported to Italy (Supplemental Figure 3). Accordingly, we classified the S segment as genotype IV. The OROV strains emerging from the Amazon region of Brazil likely arose from a new reassortment event that may contribute to the emergence of a novel genotype [1,2]. An independent analysis using the Genome Detective Virus Tool [6] confirmed that the L and S segments of C1-UF clustered within the previously described genotype IV, whereas the M segment grouped with the newly defined genotype V. A similar observation was reported by others [1], who noted that the M segment of the OROV isolated from an Italian traveler returning from Cuba did not cluster with any of the previously established genotypes.

However, because arboviruses can exist as quasispecies of variant genomes comprising major and minor virus subpopulations, we tested the hypothesis that minor OROV subpopulations can establish infections in mosquito vectors. We orally infected laboratory strains of two New World vector species, *Aedes aegypti* (Orlando, ORL) and *Anopheles albimanus* (STECLA), with OROV240023. To test the hypothesis that OROV infections in travelers may spur introduction to distant non-endemic sites, we subsequently orally infected one Old World species, *Anopheles gambiae* (KEELE), a major vector in Africa. We observed that both *Ae. aegypti* (5-10%) and *An. albimanus* (53%) established midgut infections at 7 days post-infection (7-dpi); however, we did not detect OROV 240023 in saliva (**Figure 1a-d**). In contrast, we observed that *An. gambiae* had both midgut (90%) and saliva (10%) infections at 7-dpi (**Figure 1b**). At 14-dpi, 6.6% of *Ae. aegypti* also had virus in saliva; however, *An. albimanus* never developed infectious saliva at 14 dpi. A recent study of *Aedes albopictus, Culex quinquefasciatus, Culex pipiens*, and *Anopheles quadrimaculatus* colonies indicated a lack of competence for OROV240023 saliva transmission by these vectors at 14-dpi, and of all the mosquitoes tested, only a single *Cx. pipiens* had virus-positive saliva [3]. Another study using a Brazilian OROV isolate (02.24OROV) found that mosquitoes from a lab colony of *Cx. quinquefasciatus* from Brazil, a lab colony of *Ae. aegypti* from Thailand, as well as field-derived *Ae. aegypti* from four geographically separated cities in Brazil were all refractory to virus infection at 8-dpi [4]. In contrast, a field entomological study in Porto Velho, Brazil, detected OROV RNA in three different mosquito species, including *Cx. quinquefasciatus*, suggesting that field mosquitoes may have varying vector competence for OROV [5].

**Figure 1.**
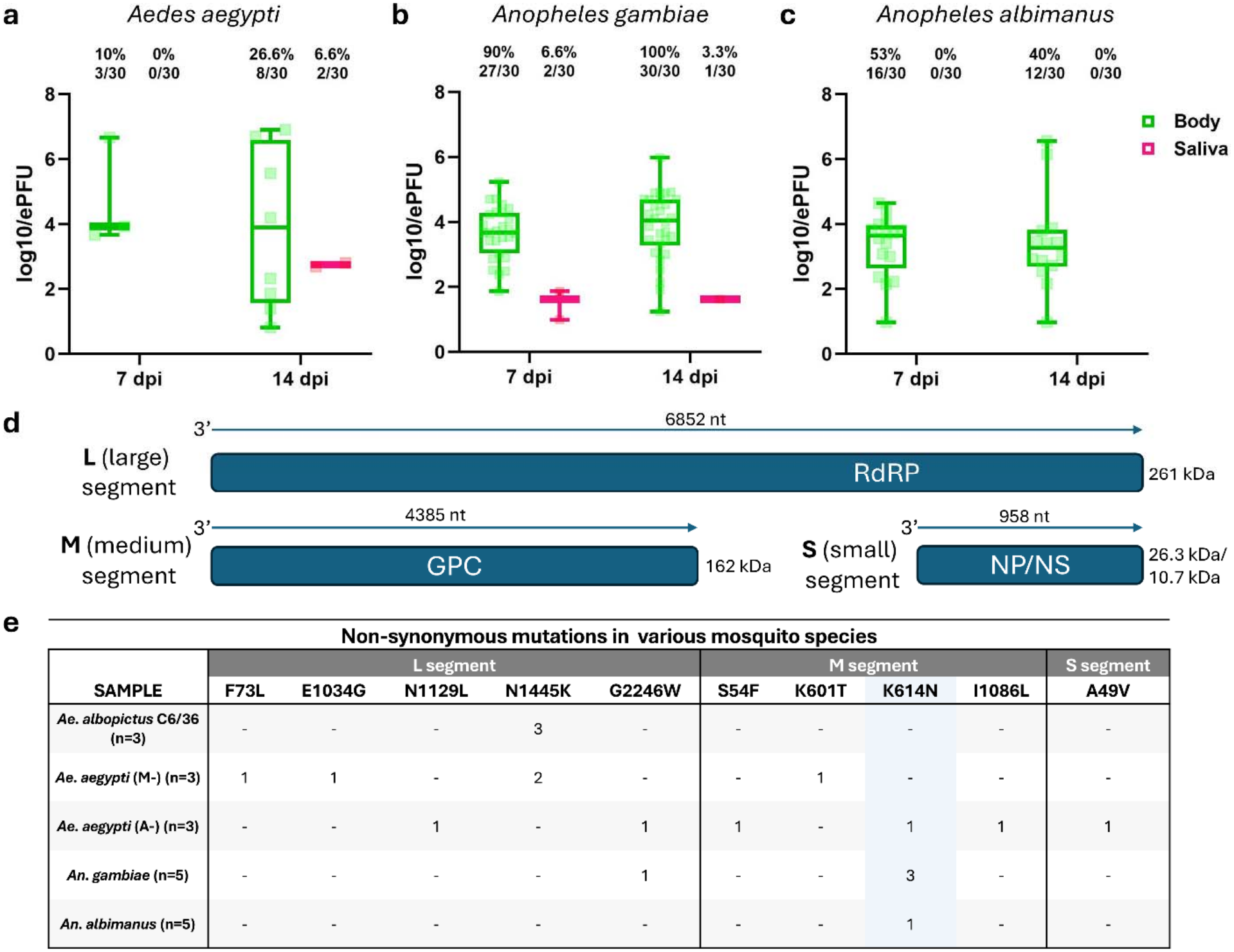
Detection of an OROV 240023 mutant that is infectious to *Aedes* and *Anopheles* mosquito vectors. Infection prevalence at 7 days and 14 days post-infection (7-dpi and 14-dpi, respectively) and estimated virus titers of OROV240023 in the body or saliva of *Aedes aegypti* (a), *Anopheles gambiae* (b), and *Anopheles albimanus* (c). The whisker-box plots highlight the median, as well as the 25^th^ and 75^th^ quartiles. OROV tripartite genome organization (d). A common non-synonymous mutation was identified across three different mosquito vectors (e). “M-” and “A-” indicate two *Ae. aegypti* mosquito cohorts.

The OROV RNA genome consists of three segments, Large (L), Medium (M), and Small (S), which encode the RNA-dependent RNA polymerase (RdRp), the glycoprotein precursor (GPC), and the nuclear (N) and non-structural proteins (NSs), respectively (**Figure 1d**) [2]. We sequenced the virus from individual infected mosquitoes to compare input and output viral genomes (**Figure 1e**). The input OROV genome from Vero E6, Aag2, and C6/36 cells matched the sequenced OROV 2024 epidemic isolates [3,7], and phylogenetic analyses grouped this isolate with other 2023-24 outbreak sequences (Supplemental Figures S1-3).

Sequencing of output genomes provided insight into the potential infectivity of OROV for different mosquito hosts. The L and M segments showed greater variability in the number of non-synonymous mutations than the S segment, which was the most conserved across all species. Only one substitution in the S segment (A49V) was detected in the nucleocapsid (N) in a single *Ae. aegypti* output genome (**Figure 1e**). Initial OROV infection revealed a K614N substitution in the M segment glycoprotein Gc, detected in 1/3 *Ae. aegypti*, 1/5 *An. albimanus*, and 3/8 *An. gambiae* abdomen + thorax sequences. Within the L segment encoding the polymerase RdRp, the N1445K substitution was detected in early *Ae. aegypti* infections and subsequently stabilized across C6/36 passages (Supplemental Table S1), suggesting that this polymerase change may be retained during cell culture. Other substitutions in the L segment (e.g., E1034G, N1129L, G2246W) and M segment (S54F, K411N, K601T, I1086L) appeared sporadically across species and were not consistently maintained (**Figure 1e**).

Sequencing of the M segment glycoprotein Gc gene from OROV grown in C6/36 cells identified position 1842 as predominantly adenine, with a consensus frequency of 99.1% across two biological replicates. A minor cytosine population was consistently detected at low frequency (0.56–0.86%) in these replicates. In Vero E6 cells, we observed a higher percentage of adenine at this position (99.8%), with non-adenine bases at frequencies below 0.1%. These low-frequency variants fall slightly above the expected range of Illumina sequencing error at Q30 and do not provide strong evidence for a true mixed population at this site, although the possibility cannot be definitively excluded. Following infection of *Anopheles gambiae* mosquitoes with virus derived from C6/36 cells, a distinct shift in frequency was observed at 7-dpi: cytosine at position 1842 became dominant (92.2%), while adenine decreased to 0.8%. This nucleotide substitution (1842:A→C) corresponds to a lysine-to-asparagine mutation at amino acid position 614 (K614N) in the Gc protein, which is involved in cell attachment.

To determine whether these observations are not random, due in part to the error-prone RdRp, which is then selected during passage through C6/36 cells and mosquitoes, we replicated the entire infection cycle using *An. gambiae*. We observed the exact K614N mutation in OROV from the saliva of an infected *An. gambiae* at 14-dpi (2.5%, 1/40), suggesting that this OROV variant already existed in the original input virus (C6/36-derived) sample as a minor population. One additional nonsynonymous mutation in the M segment glycoprotein was observed at nucleotide position 2635 (C→T), resulting in an R878C substitution in the 7-dpi *An. gambiae* OROV sequence relative to C6/36 and Vero E6 sequences. No substitutions were observed in the L and S segments, which encode the polymerase and nucleoprotein, respectively.

We noted that the reference sequence for BeAn19991 (NC_005775.1) contained the K614N mutation, whereas several other published OROV sequences (**Figure 2a**), including the TRVL9760 (KC759123.1) analyzed previously by others for mosquito infectivity [3] did not. Although it is unclear whether others have sequenced their OROV isolate for this mutation and evaluated transmissibility to the same mosquito species described herein, this observation suggests that this variant may exist as a minor species within the OROV240023 virus quasispecies. We performed a TaqMan allelic discrimination assay and identified the mutant C allele carrying the K614N mutation (**Figure 2b-c**), accounting for ∼13% of the mosaic virus population from C6/36 cells. Furthermore, deeper analysis of C6/36 sequencing data identified this allele at a frequency of 0.8% (Supplemental Table S1). Subsequent virus isolation from saliva-positive *An. gambiae* (PZ554764 [L segment];PZ554764 [M segment]; PZ545760 [S segment]) and propagation in C6/36 cells (PZ545737 [L segment]; PZ554765 [M segment]; PZ545761 [S segment]) resulted in the K614N variant as the dominant, OROV240023 consensus sequence for both samples (Supplemental Table S1).

**Figure 2.**
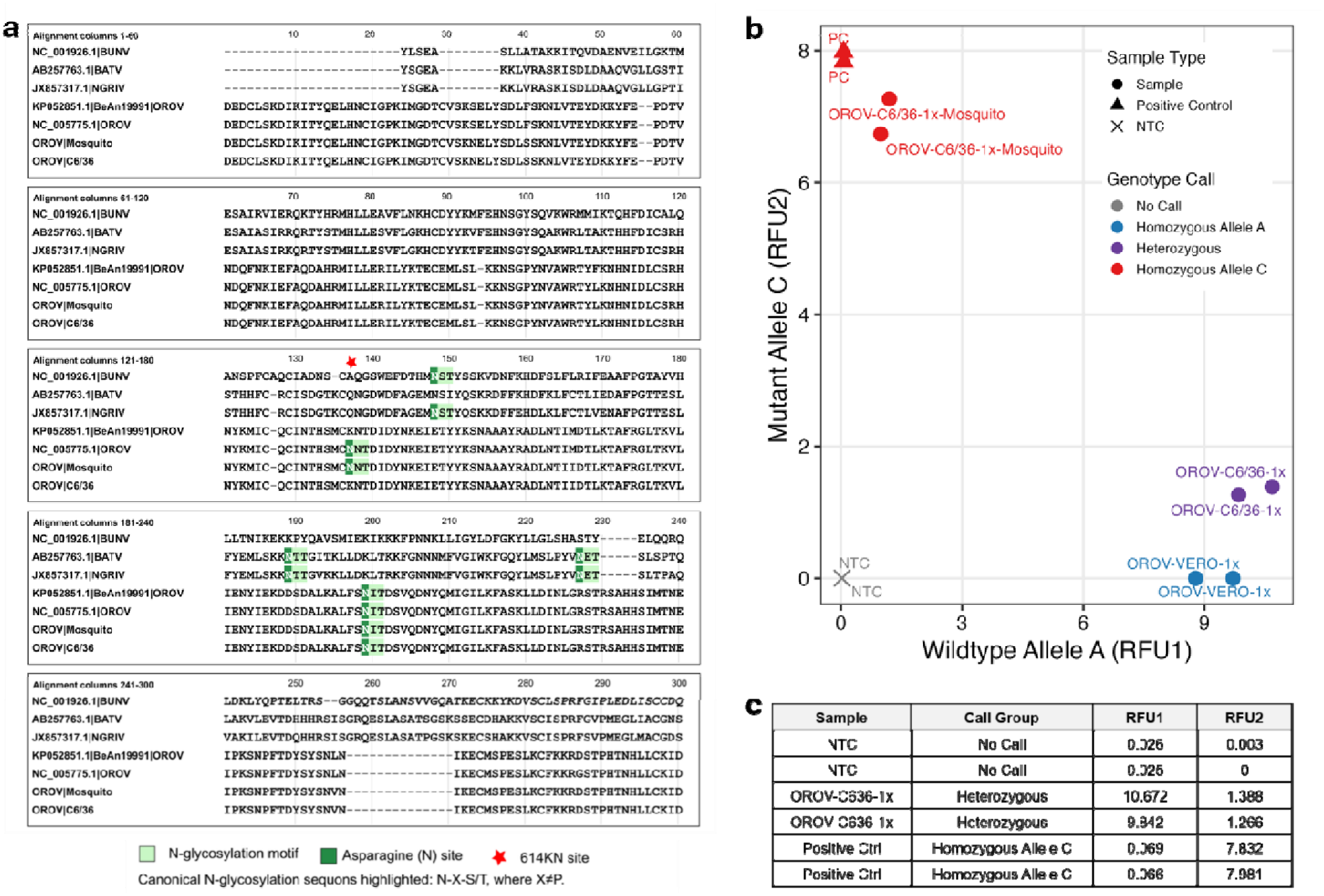
A minor variant species within the OROV240023 quasispecies is enriched in mosquito saliva. **(a)** Positional comparisons of N-glycan sites across related mosquito-borne Orthobunyavirus. **(b)** TaqMan allelic discrimination plot characterizing the proportion of wild-type Allele A and mutant Allele C associated with the K614N mutation in the Oropouche virus (OROV) Gc glycoprotein. Relative fluorescence units (RFU) for Allele A and Allele C are shown on the x- and y-axes, respectively. Samples derived from cell culture passage and/or mosquito passage were classified based on fluorescence clustering into wild-type, heterozygous, mutant, or no-call groups. A synthetic positive control carrying the mutant Allele C was included to validate assay specificity and genotype discrimination. Distinct clustering patterns clearly differentiate wild-type and mutant alleles across experimental samples. **(c)** Summary of the allelic discrimination results.

The non-synonymous substitution in the Gc glycoprotein (K164N) of the M segment (output OROV genome) from mosquitoes that had virus-positive infections resulted in a potential *N*-glycosylation site, as predicted by NetNGlyc 1.0 [8] (**Figure 2a 3a**), which differs from the BeAn19991 (prototype strain) M segment, which is identical to the original OROV240023 sequence [3] derived from the clinical sample (**Figure 3b**). A full-length model of the OROV Gc glycoprotein ectodomain (1433 amino acids) was generated using AlphaFold Server□[9] based on the sequence (**Figure 3c**). To match the overall architecture observed in published cryo-ET reconstructions of the prototypical Gc trimer [10], the model was manually trimmed in PyMOL (v2.5) [11] to retain only the well-resolved domains: head (residues 6–209), stalk (210–349), and floor (350–772). The trimmed monomer was rigid-body fitted into the subtomogram-averaged cryo-ET map (EMD-15579) using ChimeraX (v1.6) [12]. A C3-symmetric spike was reconstructed by applying 120° and 240° rotations around the Z-axis to duplicate the monomer in PyMOL, and the resulting trimeric assembly was visualized (**Figure 3d**). We generated domain-specific coloring and final structural representations in PyMOL (v2.5). These models show that the neo-glycotope, positioned relative to the stably identified *N*-glycosylation site (N676) in the parental OROV 240023, lies within an outer-facing subdomain of the tripodal spike, which may affect the cell type and species that can be infected.

**Figure 3.**
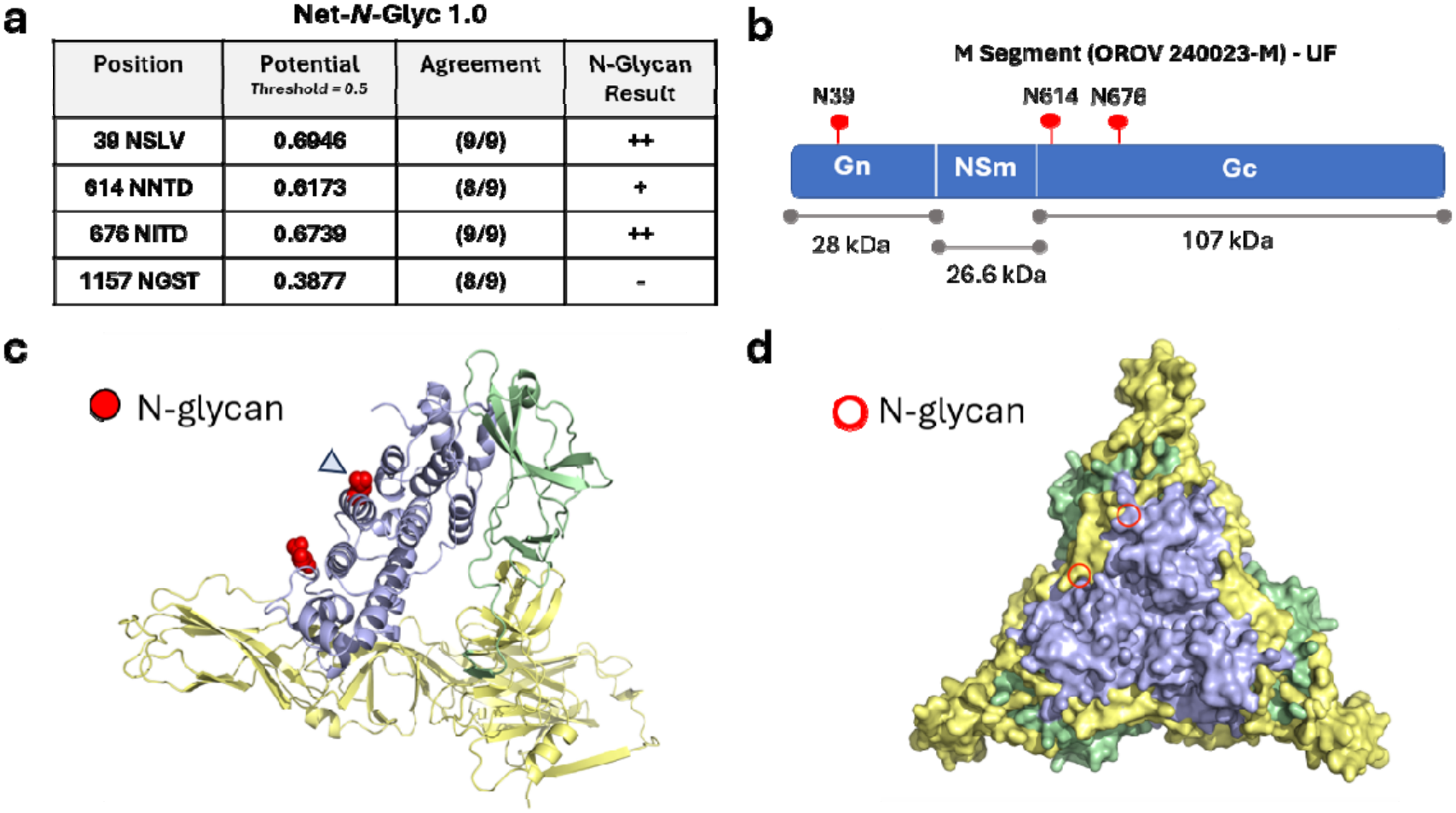
A non-synonymous substitution in the Gc glycoprotein of the M segment resulted in a new *N*-glycosylation site. **(a)** Net-*N*-Glyc 1.0 prediction of *N*-linked glycosylation sequons. **(b)** *N*-glycan positions on the M segment of OROV240023. **(c)** Ribbon model (side view) of the Gc glycoprotein and the addition of the new *N*-glycosylation site (blue arrowhead). **(d)** Space-filling model (top view) of the Gc glycoprotein demonstrating cis-positioning of the *N*-glycosylation sites.

The role of virus glycoprotein *N*-linked glycans in virus tropism has also been described for other mosquito-borne bunyaviruses [13-16], such as Bunyamwera virus [13-16] and Rift Valley Fever virus [17], pointing to C-type lectin (CTL)-mediated infection of mammalian cells [18]. It is well known that the basal lamina of mosquito salivary glands expresses CTLs, and the type and degree of expression differ across mosquito genera and species [19]. Furthermore, depending on the CTL gene family member, these lectins can either facilitate or prevent arbovirus infection of the salivary glands [19]. Our observations compel further exploration of the functional role of the second *N*-glycosylation site in OROV240023 and its possible relation to saliva-mediated infections in *Ae. aegypti* and *An. gambiae*, but not *An. albimanus*, as well as the expansion of the study to include other Culicidae.

Taken together, our observations suggest that mosquito transmission of OROV240023 cannot be completely ruled out. Although infection prevalence is relatively low in these small-scale laboratory studies, infection rates of 2-10% at higher mosquito population levels, as observed in Florida, can still effectively seed local OROV transmission. Our data demonstrate the importance of tracking OROV genomic changes across different hosts that confer functional consequences, as well as accounting for host and culture effects when evaluating OROV adaptation, selection, and transmission. However, our study is limited to laboratory colonies of mosquito vectors, and other studies have shown that vector competence for several arboviruses is also influenced by the vector host’s intrinsic microbiome and virome, as well as by interactions between virus and vector genomes [20]. Furthermore, NetNGlyc 1.0 may underestimate glycosylation sites, as BUNV has two glycosylation sites on Gc [13], yet the algorithm failed to predict glycosylation at the second site. Direct confirmation of the glycosylation sites and their mechanistic role in enabling OROV240023 infection in mosquitoes is needed. The influence of varying culture conditions across laboratories for generating the input virus in arthropod infections should not be underestimated. Future risk assessments of autochthonous transmission by local anthropophilic arthropod vectors should include a closer examination of the rapidly evolving viral genome. Risk assessment should also consider the impact of introduction to regions where transmission risk may be higher due to other competent mosquito vectors.

## METHODS

### Cell lines and Mosquito strains

We obtained cell lines from the American Type Culture Collection (ATCC, Manassas, Virginia, USA). Vero E6 (*Cercopithecus aethiops* (African green monkey) kidney cells, ATCC CRL 1586), BHK-21 cells (*Mesocricetus auratus* (Syrian golden hamster) kidney cells, ATCC CCL-10, C6/36 cells (*Aedes albopictus*) ATCC CRL-1660 and Aag2 cells (*Aedes aegypti*) were maintained as described previously [21-22]. Adult female *Ae. aegypti* ORL (Orlando), *An. albimanus* (STECLA) and *An. gambiae* (KEELE; a kind gift from P. Eggleston and H. Hurd from Keele University) mosquitoes were used for infection studies.

### Mosquito infection studies

Mosquitoes were fed blood meals containing a 2:2:1 mixture of O+ human RBCs, OROV240023, and heat-inactivated horse serum (HHIS). One group received 4.3 × 10□ PFU/mL OROV240023 (C6/36, passage 1), while another received 4.8 × 10□ UFP/mL OROV240023 (C6/36, passage 3). Saliva was collected for 30 min into capillary tubes containing 10 µL RBCs and 30 µL HHIS. Remaining tissues were collected and processed for RNA extraction as previously described [21–22].

### VirCapSeq

Total nucleic acid was reverse transcribed using random primers, followed by second-strand synthesis. Libraries were prepared with the Twist Library Preparation EF Kit 2.0 (Twist Biosciences, San Francisco, CA, USA), uniquely indexed, amplified, quantified, and pooled for capture with VirCapSeq oligonucleotide probes targeting complete vertebrate virus genomes represented in GenBank, RefSeq, EMBL, or GISAID. Captured libraries were sequenced on an Illumina NextSeq instrument [23-24]. Sequence data were processed using the Rapid Identification of Microbes (RIM) pipeline (https://rim2.gapp-cii.org/downloads/). FASTQ files underwent adapter removal and quality filtering with fastp, followed by human sequence subtraction using Kraken2.

Remaining reads were *de novo* assembled with SPAdes. Consensus OROV L, M, and S segment sequences were generated by mapping reads with Bowtie2 to GenBank references and manually inspecting alignments in Integrative Genomics Viewer for mixed populations. Sequences with <80% mapped reads were excluded from mutational and phylogenetic analyses. Sequences were aligned and trimmed to the RdRp (L), polyprotein (M), and nucleocapsid (S) coding regions and translated to identify nonsynonymous mutations relative to the clinical strain.

### Phylogenetic Analysis

All available Oropouche virus sequences in the NCBI nucleotide database as of June 14, 2024, were downloaded. We clustered sequences with CD-HIT at 95% similarity and manually selected representative sequences to capture geographic and temporal diversity. Sequences were genotyped using the Genome Detective Virus tool [25]. We included additional representatives of genotypes not captured during initial selection for phylogenetic context. We aligned sequences using MAFFT v1.5.0 in Geneious Prime v2025.1.2. To minimize alignment ambiguity from polymorphisms in untranslated regions, we trimmed sequences to open reading frames (ORFs). We then assessed recombination with RDP5 using RDP, GENECONV, BootScan, MaxChi, and SiScan. Events detected by at least two independent methods with p < 0.05 were considered credible. After removal of putative recombinants, the dataset contained 81 L-segment, 85 M-segment, and 51 S-segment genomes. Maximum-likelihood phylogenies were inferred using IQ-TREE v2.0.3 with simultaneous model selection and tree construction. Node support was assessed with 1,000 ultrafast bootstrap replicates. Trees were visualized and annotated using FigTree v1.4.4 and Inkscape v1.2.

### Allelic Discrimination Assay

RNA was reverse transcribed using the SuperScript IV First-Strand Synthesis System (Thermo Fisher Scientific) with random hexamers. We included a synthetic positive control containing the mutant allele C to validate assay specificity. A custom TaqMan SNP Genotyping Assay (ID: ANDKKXP; Applied Biosystems; Cat# 4332077) targeted the OROV M-segment SNP at nucleotide position 1,842 (A/C), corresponding to residues 614K/614N. VIC- and FAM-labeled MGB probes distinguished the adenine and cytosine alleles, respectively. Reactions (10 µL) contained 5 µL TaqMan Genotyping Master Mix, 0.5 µL 20× assay mix, and 4.5 µL cDNA template (15 ng). Cycling consisted of 95°C for 20 s, followed by 40 cycles of 95°C for 3 s and 60°C for 30 s. Allelic discrimination was performed using Azure Cielo Real-Time PCR System software, and genotypes were assigned by clustering relative fluorescence units (RFU) as homozygous wild-type, homozygous mutant, or heterozygous. We exported and plotted endpoint FAM and VIC RFU data in R v4.0.0 using ggplot2 and ggrepel.

### Structural Modeling

The OROV240023 Gc model was generated using the AlphaFold Server from the Gc sequence derived from the experimentally determined M-segment sequence. We generated five models and selected the highest-scoring model based on pLDDT and predicted aligned error. Low-confidence unstructured termini and loops were removed in PyMOL v2.5, retaining the head (residues 6–209), stalk (210–349), and floor (350–772) regions. Residue assignments were cross-checked against available orthobunyavirus Gc head and stalk structures.

*N*-linked glycosylation sites were predicted from the complete M-segment precursor using NetNGlyc 1.0 with default settings. Sites matching the N-X-S/T consensus (X ≠ Pro) with positive prediction scores were retained, and N133 and N195 were mapped onto the AlphaFold model.

For structural fitting, the ∼16 Å subtomogram-averaged Bunyamwera virus spike density (EMD-15579) was fitted with the trimmed OROV Gc monomer in UCSF ChimeraX v1.6. Multiple initial placements were tested, and the fit was evaluated by correlation coefficient. A tripodal spike was reconstructed in PyMOL by generating two additional copies of the fitted monomer and rotating them by 120° and 240° about the threefold axis. Final structural figures were prepared using PyMOL and ChimeraX.

## Data Availability

All virus sequences are available in GenBank using the accession numbers provided in Supplemental Table S1.

## Author contributions

L.A.A.P., J.C.P., B.A., X.J.L., D.L., and R.R.D. designed the experiments, analyzed results, and wrote the paper. T.B., J.A.L., N.L.T., and J.K.W. analyzed the results, wrote and reviewed the paper. R.R.D. conceived the study and provided overall supervision.

