## Supplemental Figures for "An Oropouche virus *N*-glycosylation mutation in the Gc protein head domain is associated with transmissible infection in *Anopheles* mosquitoes"

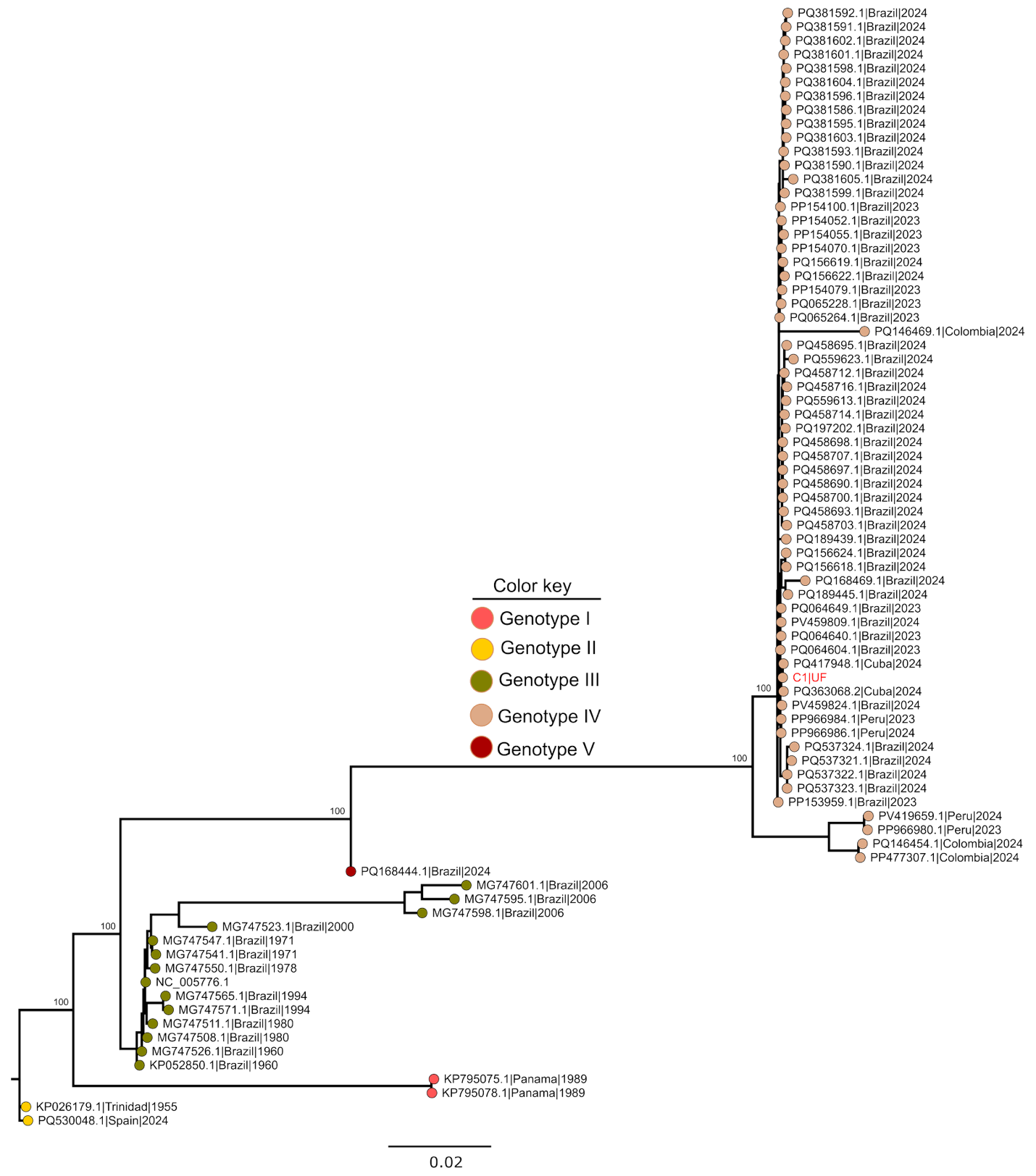

**Figure S1. Maximum likelihood phylogeny of a subset of global Oropouche virus genomes (L-segment).** The phylogeny was constructed from the complete coding sequence of the L-segment (UF OROV strain, red text). Strains are labeled with their NCBI accession numbers, country of origin, and collection date. Node numbers represent bootstrap values, and scale represents the number of nucleotide substitutions per site. The phylogeny was tested with 1000 bootstrap replicates.

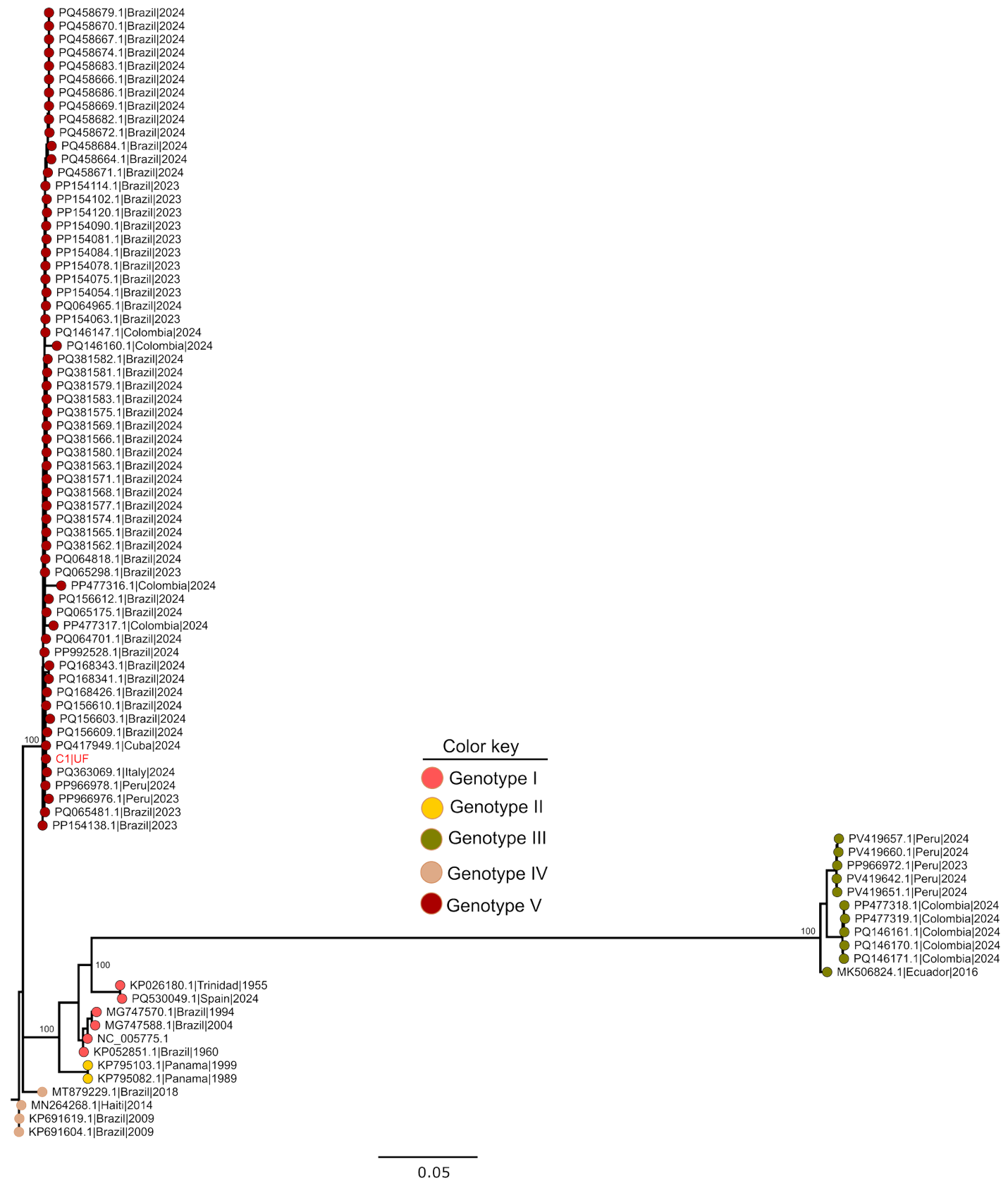

**Figure S2. Maximum likelihood phylogeny of a subset of global Oropouche virus genomes (M-segment).** The phylogeny was constructed from the complete coding sequence of the M-segment (UF OROV strain, red text). Strains are labeled with their NCBI accession numbers, country of origin, and collection date. Node numbers represent bootstrap values, and scale represents the number of nucleotide substitutions per site. The phylogeny was tested with 1000 bootstrap replicates. The genotype switch reflects OROV240023 as a novel reassortant with significant changes in the M segment compared to historical strains.

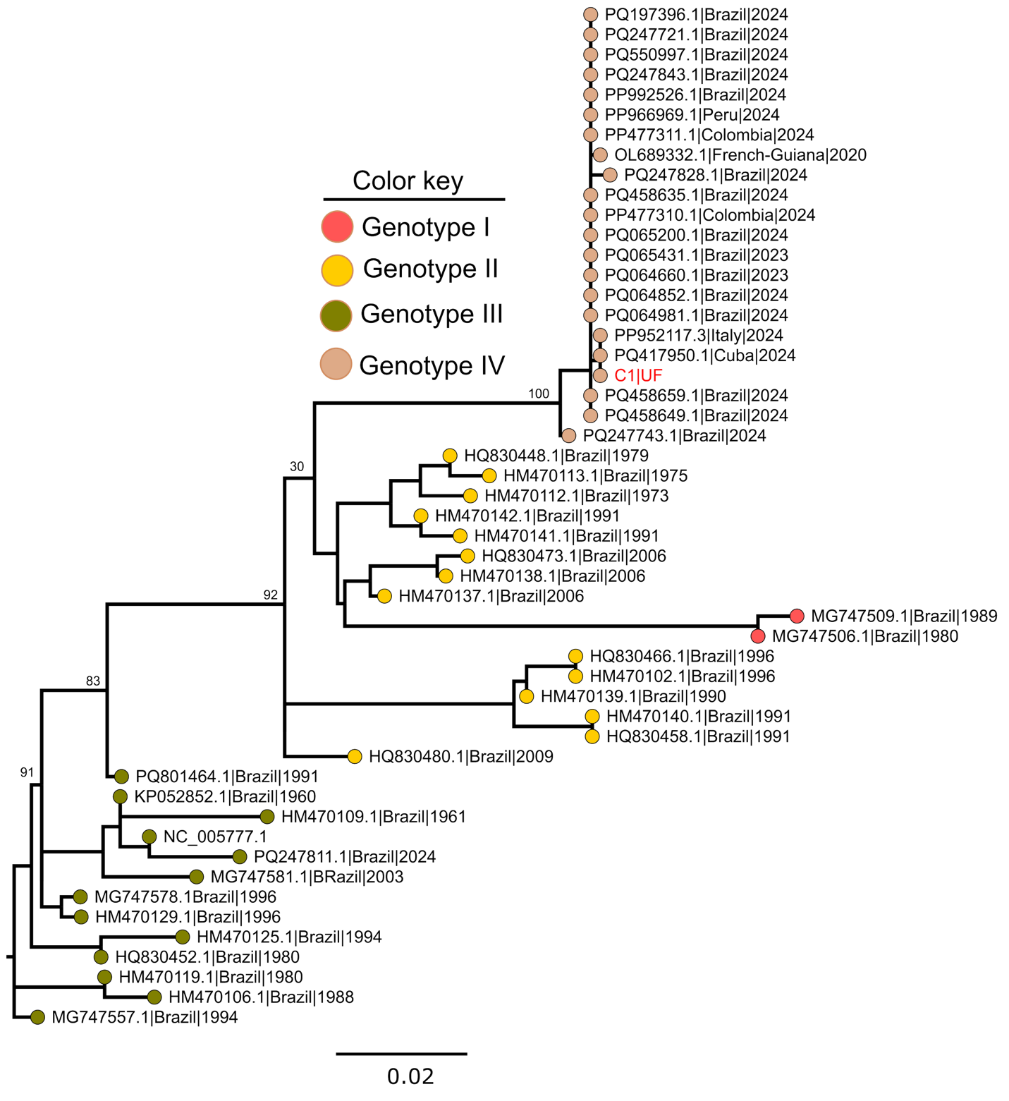

**Figure S3. Maximum likelihood phylogeny of a subset of global Oropouche virus genomes (S-segment).** The phylogeny was constructed from the complete coding sequence of the S-segment (UF OROV strain, red text). Strains are labeled with their NCBI accession numbers, country of origin, and collection date. Node numbers represent bootstrap values, and scale represents the number of nucleotide substitutions per site. The phylogeny was tested with 1000 bootstrap replicates.
